# Cold Quad-Modal Nanocomplex for Precise and Quantitative *in Vivo* Stem Cell Tracking

**DOI:** 10.1101/2025.04.16.649154

**Authors:** Ali Shakeri-Zadeh, Chao Wang, Shreyas Kuddannaya, Saleem Yousf, J.W.M. Bulte

**Affiliations:** The Russell H. Morgan Department of Radiology and Radiological Science, Division of MR Research, The Johns Hopkins University School of Medicine, Baltimore, MD, USA; Cellular Imaging Section and Vascular Biology Program, Institute for Cell Engineering, The Johns Hopkins University School of Medicine, Baltimore, MD, USA; The Russell H Morgan Department of Radiology and Radiological Science, Division of Cancer Imaging Research, The Johns Hopkins University School of Medicine, Baltimore, MD, USA; Department of Biomedical Engineering, The Johns Hopkins University School of Medicine, Baltimore, MD, USA; Department of Oncology, The Johns Hopkins University School of Medicine, Baltimore, MD, USA; Department of Chemical & Biomolecular Engineering, The Johns Hopkins University Whiting School of Engineering, Baltimore, MD, USA; F.M. Kirby Research Center for Functional Brain Imaging, Kennedy Krieger Inc. Baltimore, MD, USA

## Abstract

Current single imaging modalities typically lack the ability to simultaneously offer detailed anatomical visualization and quantitative cellular information, which is crucial for evaluating and improving therapeutic efficacy. We developed a quad-modal imaging nanocomplex for magnetic resonance imaging (MRI), magnetic particle imaging (MPI), computed tomography (CT), and multispectral optoacoustic tomography (MSOT) within a single nanoplatform. The chemically engineered complex is composed of bovine serum albumin as biocompatible matrix, superparamagnetic iron oxide as MRI and MPI agents, and optoradiopaque bismuth sulfide as CT and MSOT agents. We demonstrate here its use for high-resolution, real-time, and quantitative *in vivo* imaging of mesenchymal stem cells transplanted in mouse brain. This versatile nanocomplex may find applications for monitoring cell transfer and cell transplantation *in vivo* using multiple imaging approaches.

## Main

Further advances in stem cell therapy and regenerative medicine will benefit from *in vivo* imaging techniques for cell localization and quantification^1–4^. Numerous approaches for *in vivo* tracking of stem cells have been used in preclinical and clinical studies, including the use of positron emission tomography (PET)^5^, single photon emission tomography (SPECT)^6^, bioluminescence imaging (BLI)^7^, computed tomography (CT)^8^, magnetic resonance imaging (MRI)^9,10^, magnetic particle imaging (MPI)^11,12^, and multispectral optoacoustic tomography (MSOT)^8^. Each imaging modality has its own strengths and weaknesses for tracking cells *in vivo* (see Table 1).

**Table 1:** Comparison of the different clinical imaging modalities that are available for *in vivo* tracking of stem cells. **Red**: Modalities that use “hot” tracers. **Blue:** Modalities that use “cold” tracers or contrast agents used as quad-modal agents in this study.

| Criteria | PET | SPECT | US | CT | MRI | MPI | MSOT |
| --- | --- | --- | --- | --- | --- | --- | --- |
| Ionizing radiation | Yes | Yes | No | Yes | No | No | No |
| Sensitivity | Excellent | Excellent | Excellent | Limited | Moderate | Excellent | Excellent |
| Specificity | Excellent | Excellent | Excellent | Limited | Limited | Excellent | Moderate |
| Quantification | Moderate | Moderate | Limited | Excellent | Limited | Excellent | Moderate |
| Anatomical content | No | No | Moderate | Excellent | Excellent | No | Moderate |
| Spatial resolution | Limited | Limited | Moderate | Excellent | Excellent | Moderate | Moderate |
| Temporal resolution | Limited | Limited | Excellent | Excellent | Excellent | Excellent or Moderate | Excellent |

Multimodal imaging may be used to combine strengths and obviate weaknesses when there is a simultaneous, complementary reporting on cell biodistribution, quantity and viability with high resolution, specificity, and sensitivity^13^. The effective use of multimodal imaging hinges on specific clinical or preclinical aims, and is dependent on multiple single imaging agents or ideally a multimodal imaging probe including those based on nanoparticles (NPs)^14^. Hence, further development of versatile NPs that can be detected with multiple imaging modalities is warranted^15^. Several studies have developed tri-modal^16,17^ and quad-modal^18,19^ agents to enhance the imaging information across multiple modalities. However, most reported multimodal NPs rely on incorporating “hot (radioactive)” agents for quantitative tracking using PET and SPECT^18–20^, and require bespoke manufacturing and radiation safety processes.

Advancements in MPI and MSOT technologies have recently helped overcome such limitations. MPI is sensitive, specific, and quantitative^11^, but it lacks anatomical information and an excellent spatial resolution for cell tracking, while MSOT offers quantification^21^,spectral selectivity and high spatiotemporal resolution^22^. Hence, the development of “cold” multimodal NPs that respond to MRI and CT as anatomical imaging modalities, and MPI and MSOT as quantitative methods will offer opportunities to paint a comprehensive picture of *in vivo* cell distribution.

We report here on a “cold” versatile nanocomplex for quad-modal cell tracking that combines superparamagnetic iron oxide (SPIO) NPs and optoradiopaque bismuth sulfide (Bi_2_S_3_) NPs within a bovine serum albumin (BSA) matrix. This integration leverages the magnetic properties of SPIO for MRI^10^ and MPI^11^, with the dual optical and radiopaque characteristics of Bi_2_S_3_ providing contrast for CT^23^ and MSOT^24^. The BSA matrix enhances the biocompatibility and stability of the nanocomplex^25^, ensuring its integrity as a single platform suitable for comprehensive multimodal imaging.

### Nanocomplex synthesis and physicochemical characterization

Using a step-by-step solvothermal decomposition method, **<u>AB</u>**nanocomplexes composed of **<u>A</u>**lbumin and **<u>B</u>**i_2_S_3_, were synthesized first, followed by the addition of **<u>S</u>**PIO to form **ABS**. Ferucarbotran (a carboxydextran-coated iron oxide and the active pharmaceutical ingredient in Resovist®) was chosen as SPIO for the ABS nanocomplex due to its commercial availability (ensuring quality control and widespread use) and excellent properties for both MRI^26^ and MPI^27,28^. While Resovist^®^ was discontinued by Bayer-Schering Pharma (Berlin, Germany), ferucarbotran is still manufactured in Japan by Meito-Sangyo Co. Ldt. (Nagoya, Japan), relabeled and diluted by Magnetic Insight Inc. as Vivotrax^®^. In addition, Bayer-Schering has now rebranded Resovist^®^ as Resotran^29^, aimed at its use as clinical MPI tracer. Bi_2_S_3_ is a better opto-radiopaque agent than the more commonly used gold NPs, as it shows a narrower bandgap and a larger X-ray attenuation coefficient (1.3 vs. 5 eV and 5.8 vs. 5.1 cm^−2^kg^−1^ at 100 KeV, for Bi_2_S_3_ and gold NPs, respectively)^30^.

The chemical composition, functional groups, and optical properties of ABS were characterized by Fourier transform infrared (FTIR) and ultraviolet-visible (UV-Vis) spectroscopy. Surface charge, size, particle concentration, morphology, composition, and protein binding assessments of ABS were performed by zeta potential analysis, dynamic light scattering (DLS), nanoparticle tracking analysis (NTA), transmission electron microscopy (TEM), high-angle annular dark field (HAADF), inductively coupled plasma mass spectrometry (ICP-MS), multi-angle light scattering (MALS), and asymmetric-flow field-flow fractionation (AF4). Key findings are summarized in **Fig.1**, while complementary characterization results are provided in the Supplementary Information (**Figs. S1-S6 and Tables S1-S11)**

**Figure 1:**
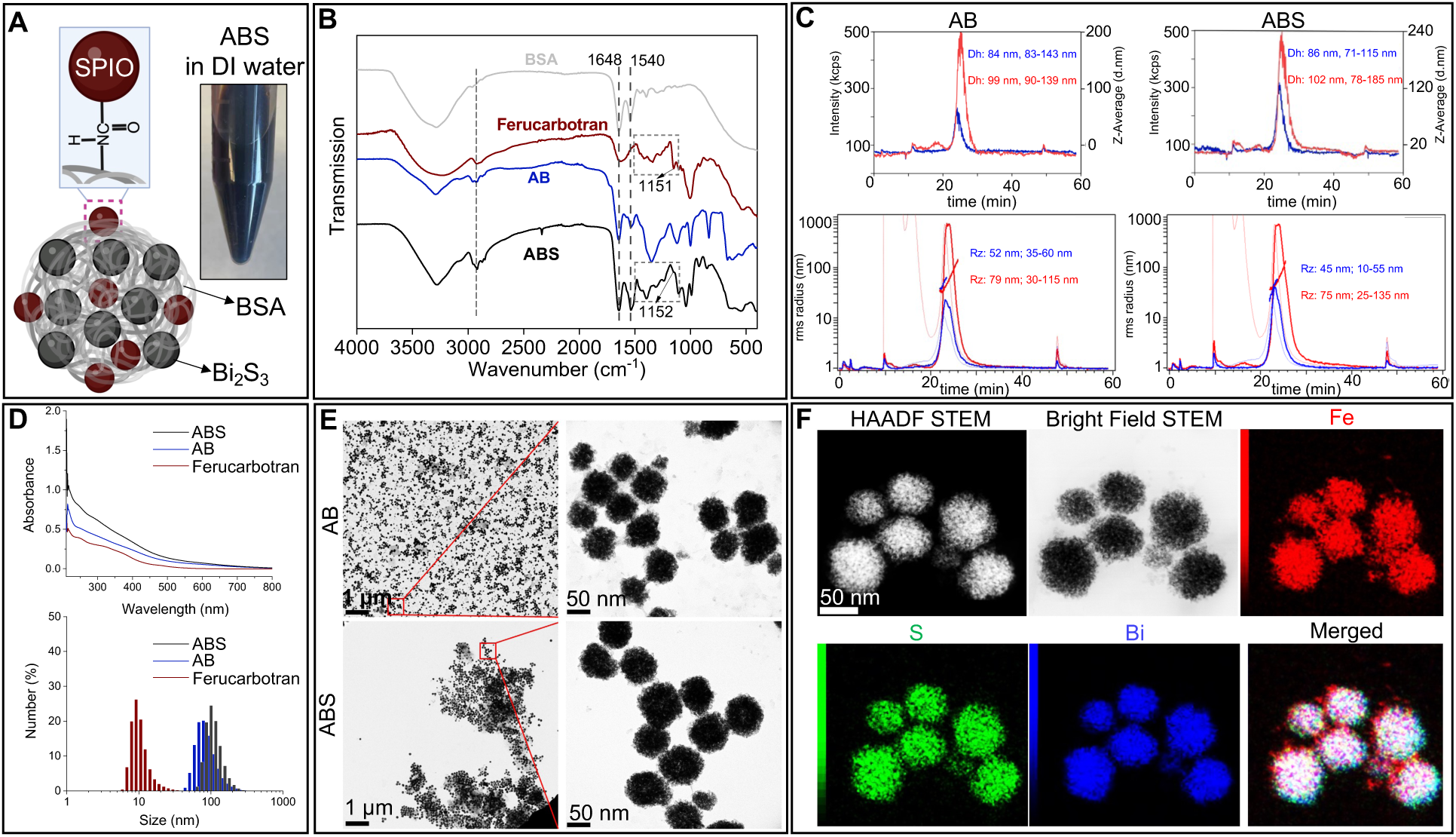
Physicochemical characterization of ABS. (**A**) Schematic illustration and appearance of ABS. (**B**) FTIR spectra comparing BSA, ferucarbotran, AB, and ABS. (**C**) Particle size distribution for AB and ABS measured using AF4 coupled with MALS and DLS detectors to determine protein binding to their surface. (**D**) UV-Vis absorbance spectra and size distribution histograms of ferucarbotran, AB, and ABS. (**E**) TEM images of AB and ABS at different magnifications. (**F**) HAADF STEM and bright-field STEM images and elemental mapping showing the nanoscopic distribution profile of Fe, S, and Bi.

**Fig.1A** depicts the structure and appearance of ABS in DI water as a dark near-black formulation. FTIR spectra show the BSA peaks at 1648 cm^-1^ and 1540 cm^-1^, characteristic of amide I (C=O stretching vibrations) and amide II bands (N-H bending and C-N stretching), respectively (**Fig. 1B**). The peak in the BSA spectrum just below 3000 cm^-1^ represents CH_2_ stretching vibrations. These vibrations across the spectra of BSA, AB, and ABS indicates that BSA is indeed integrated with the AB and ABS, and that the addition of SPIO and Bi_2_S_3_ NPs does not significantly alter the overall structure of the BSA matrix. In the ferucarbotran and ABS spectra, the Fe-O stretching observed between 600–550 cm^-1^ corresponds to the SPIO core. The spectral peaks around 3400 cm^-1^ for O-H stretching, 1750–1700 cm^-1^ for C=O stretching, and 1150 cm^-1^ for C-O stretching are indicative of the carboxydextran coating of the SPIO core.

Particle size distributions for AB and ABS were measured using AF4 coupled with MALS and DLS detectors. Samples were also incubated with human plasma to assess surface protein binding. AB had one major peak with an averaged hydrodynamic diameter of 84 nm which increased to 99 nm after plasma incubation (**Fig. 1C**). The corresponding radius of gyration sizes for AB were 52 and 79 nm, respectively. This resulted in a shape factor (2*Rg/Dh) of 1.23 without plasma incubation and 1.59 after plasma incubation, implying protein binding to the surface of the AB. Similar to AB, ABS also had one major peak with an averaged hydrodynamic diameter of 86 nm. Following plasma incubation, the size increased to 102 nm (**Fig. 1C**). The corresponding radius of gyration sizes for ABS were 45 and 75 nm, respectively, resulting in a shape factor of 1.04 without plasma and 1.47 after plasma incubation, demonstrating that there was a similar protein binding to the surface of ABS.

UV-Vis spectra demonstrate that the absorbance of ferucarbotran decreases with increasing wavelength, a feature typical for metal oxide NPs^31^ (**Fig. 1D**). However, for AB and ABS which incorporate metal sulfide NPs, different optical properties exist due to their unique electronic structures and different light interactions^32^. The variations in the absorption profiles for ABS compared to AB and ferucarbotran likely result from the combined effects of SPIO and Bi_2_S_3_, illustrating the complex interaction of these components within the ABS nanocomplex. DLS analysis revealed that ferucarbotran exhibits a peak particle size of 8.5 nm and a Z-average of 40.6 nm (**Fig. 1D**). AB displays a peak size of 75.6 nm with a Z-average of 104.3 nm, while ABS shows a peak size of 104.5 nm with a Z-average of 135.1 nm. Further characterization data on particle sizing, concentration assessment, and quantitative elemental analysis are provided in the Supplementary Information (**Figs. S1-S5, Tables S1-S11**).

TEM images indicate that both AB and ABS possess a spherical morphology (**Fig. 1E**). ABS is characterized by larger clusters, likely arising from the incorporation of SPIO. ABS was also characterized using HAADF STEM, bright field STEM, and elemental analysis to determine its morphology and composition at high-definition (**Fig. 1F**). The findings demonstrate the integration of multi-component NPs within the ABS formulation, highlighting a uniform physical structure/size and good dispersibility, as well as the effective encapsulation and distribution of SPIO and Bi_2_S_3_ NPs within the BSA.

Full zeta potential analysis of AB and ABS at native pH of 7.29 showed negative values of-23.1±1.5 and-24.3±1.0 mV, respectively, while ferucarbotran was neutral at native pH of 7.29 (**Fig. S6, Table S11**. The negative zeta potential for ABS suggests that the dextran coating is possibly interweaved within the BSA matrix.

To enhance the cell detection sensitivity of CT to that of MRI and MPI, which are inherently more sensitive, we designed ABS with an initial Bi to Fe mass ratio of 5:1.

This ratio was intended to enhance CT detectability when the same probe is used for cell labeling. The actual concentrations of Bi and Fe in ABS were quantified using ICP-MS, confirming an experimental ratio aligning with the theoretical ratio (**Table S8**).

### Quantitative *in vitro* imaging studies of naked ABS

The imaging properties of ferucarbotran, AB, and ABS are shown in **Fig. 2**.

**Figure 2:**
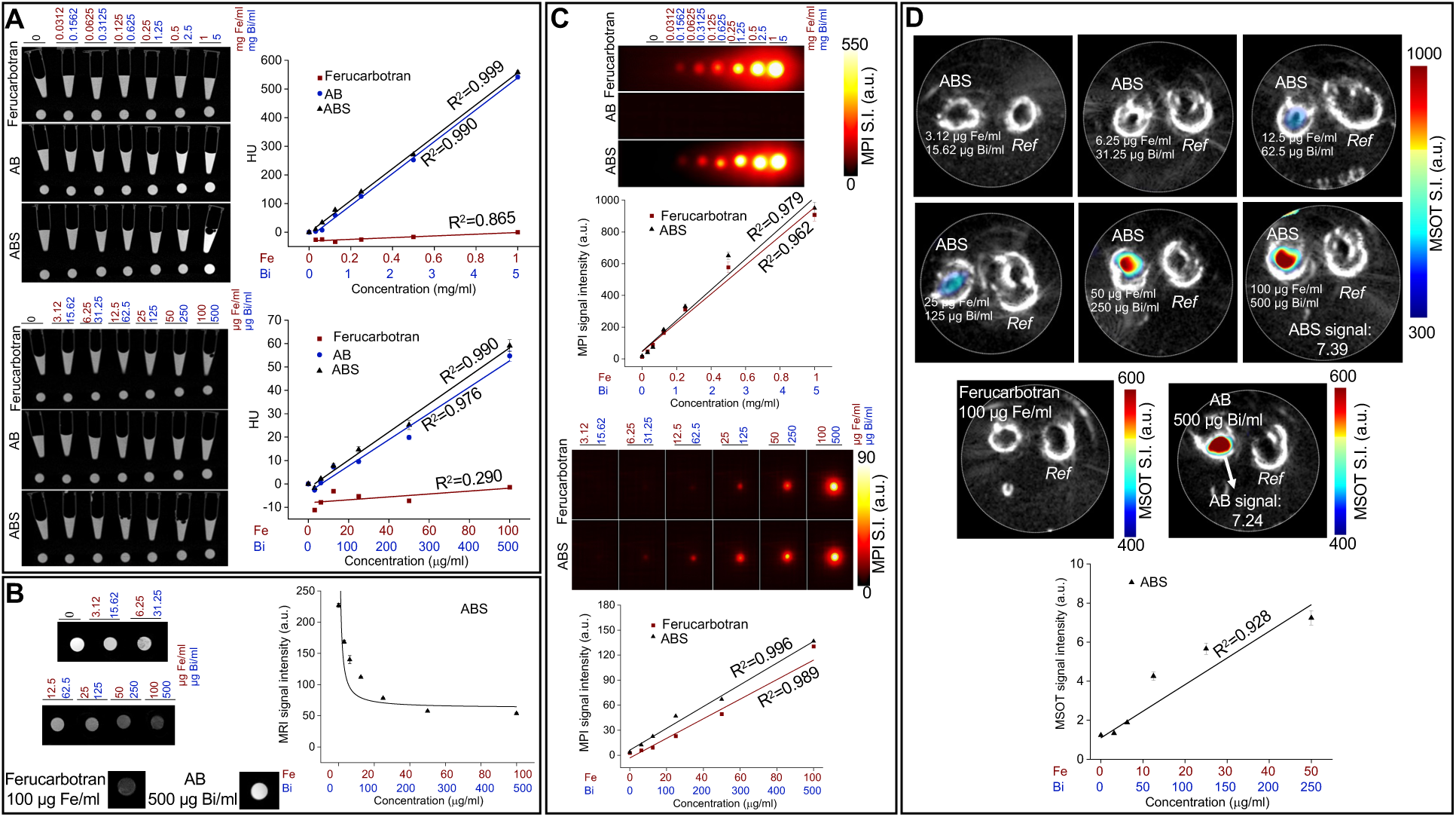
**Quantitative phantom imaging studies**. Two different concentration categories, including high (1 mg Fe/ml or 5 mg Bi/ml) and low (100 µg Fe/ml or 500 µg Bi/ml) concentrations were tested. (**A**) CT imaging of serial dilutions of ferucarbotran, AB, and ABS formulations with corresponding HU values versus concentration. (**B**) MRI and signal intensity analysis of ABS. The highest equivalent concentration per Fe for ferucarbotran and AB were also imaged with MRI under the same conditions and are shown as the control group for ABS. (**C**) MPI comparing different concentrations of ferucarbotran, AB, and ABS formulations. (**D**) MSOT of ABS, ferucarbotran, and AB formulations. The highest equivalent concentration per Bi for ferucarbotran and AB were also imaged by MSOT and shown as the control group for ABS. “Ref” in panel **D** contained 100 µl of DI water. Data in each panel represent three independent experiments.

Quantification of CT contrast in Hounsfield Units (HU) revealed no contrast enhancement for ferucarbotran, but AB and ABS demonstrated a linear increase in CT contrast with concentration (R^2^=0.98-1.00, **Fig 2A**). T2-weighted (w) MRI showed that there is no such relation in signal intensity for ABS, inherent to the non-quantitative nature of MRI for detecting SPIO. We observed no differences in MRI signal intensity between ABS and ferucarbotran for the same Fe concentration, nor did AB show notable changes compared to water at equivalent Bi concentrations. As expected, AB was undetectable by MPI, whereas ABS and ferucarbotran exhibited a strong MPI signal (**Fig. 2C**). The MPI performance of ABS was slightly higher than ferucarbotran, suggesting underlying differences in magnetic relaxation that warrant further physicochemical investigation. MSOT also proved to be quantitative, as evidenced by the linear response (R=0.93) of signal vs. concentration (**Fig. 2D**). No differences in MSOT signal intensity were observed between ABS and AB at equivalent Bi concentrations. Similarly, ferucarbotran at the same Fe concentration as ABS showed no change in MSOT signal intensity compared to water.

Thus, except for MRI, the amount of ABS can be quantified with CT, MPI, and MSOT as they exhibit a linear response with concentration. The minimum detectable ABS concentration for MSOT was approximately 12.5 µg Fe/ml or 62.5 µg Bi/ml (**Fig. 2D**), which is 2 times higher than that for MPI (**Fig. 2C**). Given a Bi:Fe ratio of 5:1, this suggests that the MPI sensitivity for Fe detection is effectively 10 times higher than the MSOT sensitivity for Bi detection. Based on clinical protocols, an HU of greater than 50 is considered an acceptable minimum for contrast enhancement of CT imaging^33^, the minimum detectable ABS concentration can be considered at 100 µg Fe/ml or 500 µg Bi/ml (**Fig. 2A**). Consequently, the CT detection sensitivity is 8 times lower than MSOT for Bi detection.

### Cellular uptake, cytotoxicity, and *in vitro* imaging of ABS-hMSCs

We have chosen hMSCs as a cellular prototype for ABS labeling, reflecting their extensive clinical use for treating a wide range of diseases including ischemic heart disease^34^, osteoarthritis^35^, diabetes mellitus^36^, neurological disorders^37^, and cancer^38^, with over 1,000 registered clinical trials highlighting their therapeutic relevance. **Fig. 3A** illustrates the cellular uptake and peri-nuclear localization of ABS, its parental components, ferucarbotran and AB after Prussian Blue (PB) staining. TEM was used to further assess labeling at the ultrastructural level (**Fig. 3B**). A Ferrozine spectrophotometrical assay for iron quantification revealed an average intracellular content of 92.95±6.72 pg of Fe per single cell.

**Figure 3:**
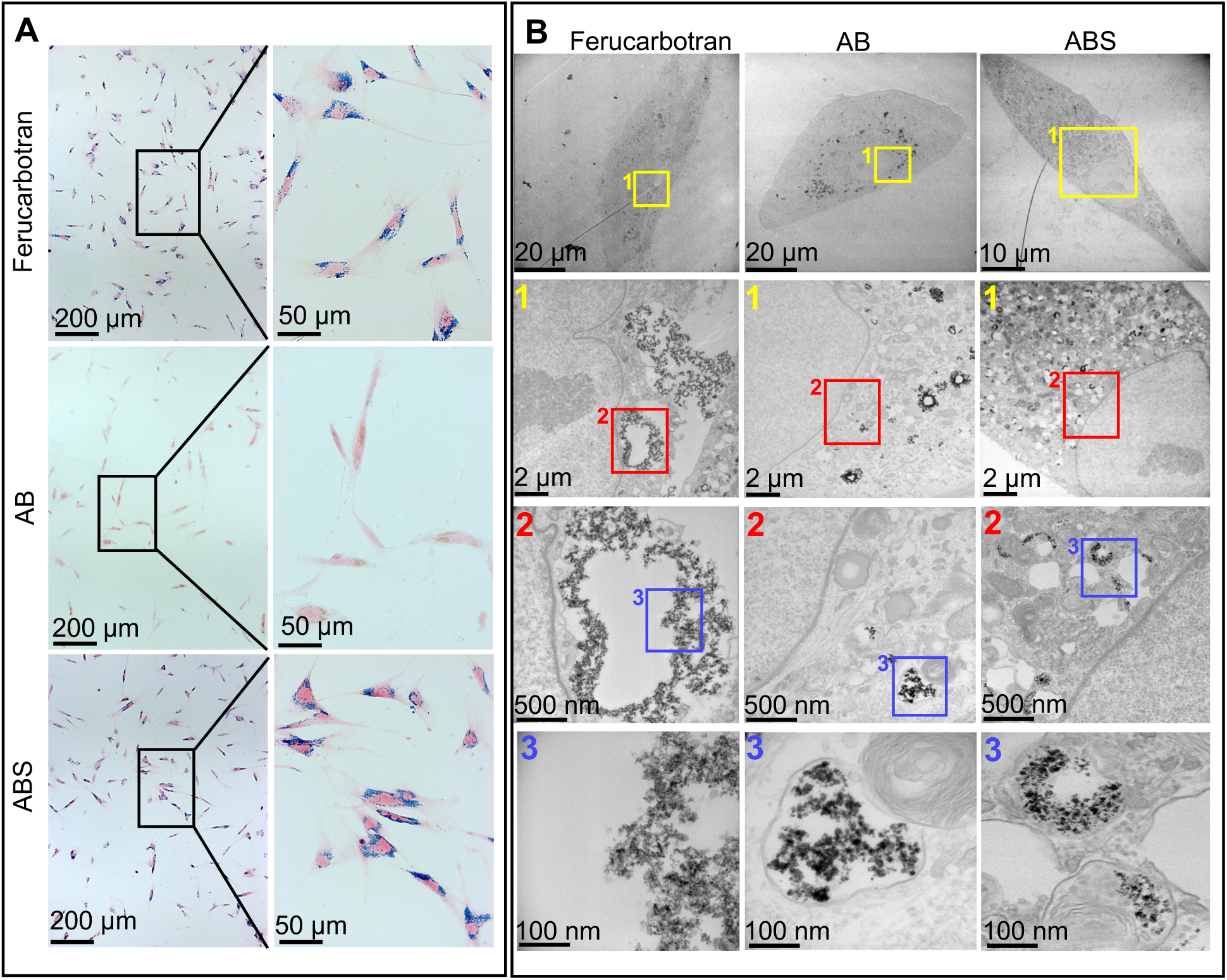
Cellular uptake of ABS. (**A**) PB staining of hMSCs labeled for 24 hrs with 25 µg Fe/ml ABS and its parental components ferucarbotran and AB. (**B**) TEM images of cells labeled with ferucarbotran, AB, and ABS at various magnifications, showing the intracellular distribution and aggregation of NPs within subcellular structures.

To evaluate the absence of cytotoxicity following labeling, hMSCs were incubated with different concentrations of ferucarbotran, AB, and ABS formulations for 24 or 72 hrs. While ferucarbotran exhibited no significant cytotoxic effects across all tested concentrations and time points, ABS started to display cytotoxicity at concentrations ≥150 µg Bi/ml after 24 hrs and ≥100 µg Bi/ml after 72 hrs incubation (**Fig. 4A**). We therefor selected a concentration of 25 µg Fe/ml equivalent to 125 µg Bi/ml and 24 hrs as the concentration and incubation time for all further experiments. hMSCs were labeled with ABS and imaged at two different cell densities, 10,000 cells/µl and 1000 cells/µl. Imaging was performed using CT (**Fig. 4B**), MPI and MRI (**Fig. 4C)**, and MSOT (**Fig. 4D**). Unlike MRI, the quantitative nature of CT, MPI, and MSOT was affirmed when R² values exceeded 0.9. CT contrast did not notably enhance at cell densities below1000 cells/µl, where HU was lower than 50. Both MPI and MSOT successfully detected a minimum cell density of ∼62 ABS-labeled hMSCs per µl. This does however not imply that MPI and MSOT have the same sensitivity for cell detection, given that the Bi:Fe ratio used was 5:1. Hence, MPI exhibits the highest cell detection sensitivity among the four imaging modalities.

**Figure 4:**
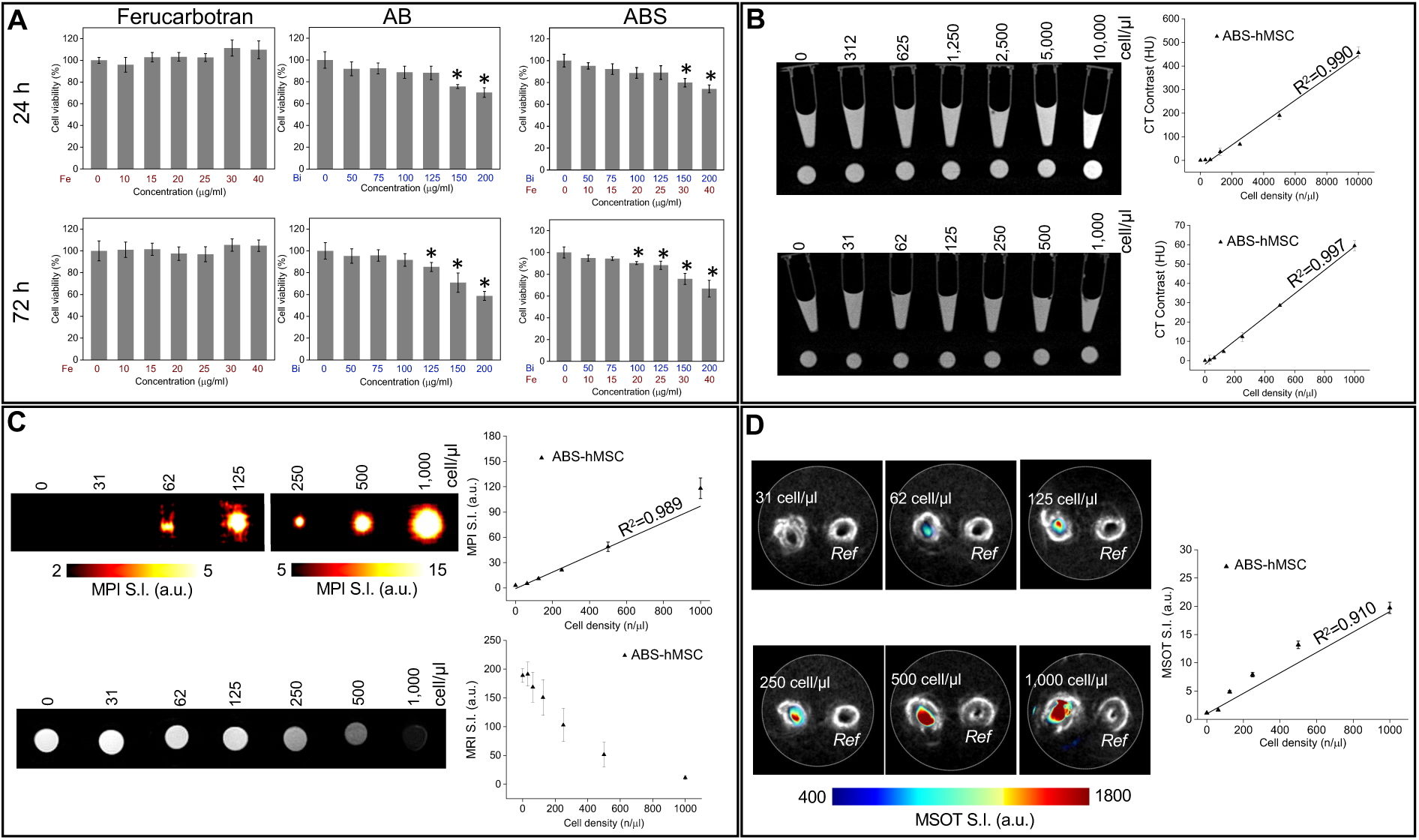
Cell viability and phantom imaging studies of ABS-labeled hMSCs. Two different cell densities were tested, 10,000 cells/µl and 1,000 cells/µl. (**A**) Cell viability of ABS-labeled hMSCs for varying concentrations of Fe or Bi and incubation times. *p<0.05 compared to control (0, unlabeled) group. (**B**) CT, (**C**) MPI/MRI, and (**D**) MSOT of serial dilutions of ABS-labeled hMSCs with corresponding quantification and linear regression analysis. “Ref” in panel **D** contained 100 µl of 8% gelatin with 0.02% w/v sodium azide added. Data in each panel figure represent three independent experiments.

### *In vivo* imaging of ABS-labeled hMSCs transplanted in mouse brain

Immunodeficient Rag2^-/-^ mice were used to prevent immediate immunorejection of transplanted hMSCs. **Fig. 5A** shows bi-modal *in vivo* MPI and MRI conducted at various time points after cell injection depicting the ability to track the spatial localization and persistence of ABS-hMSCs over time simultaneously. Both magnetic imaging modalities did exhibit blooming artifacts, with its actual size not notably changing with varying cell densities on MRI. In contrast, the size decreased with reduced cell density on MPI. *Ex vivo* CT and MSOT imaging were also conducted one-month post-injection, accompanied by histological analysis (**Fig. 5A**) using PB staining and anti-human nuclear antigen (HuNA) staining to validate the imaging findings, with a good agreement with the imaging data. All four imaging modalities were able to detect cell densities as low as 6,250 ABS-hMSCs per µl. The observation for CT, despite being the least sensitive method tested, is in agreement with the *in vitro* detection threshold being 1,000 ABS-hMSCs per µl (**Fig. 4B**).

**Figure 5:**
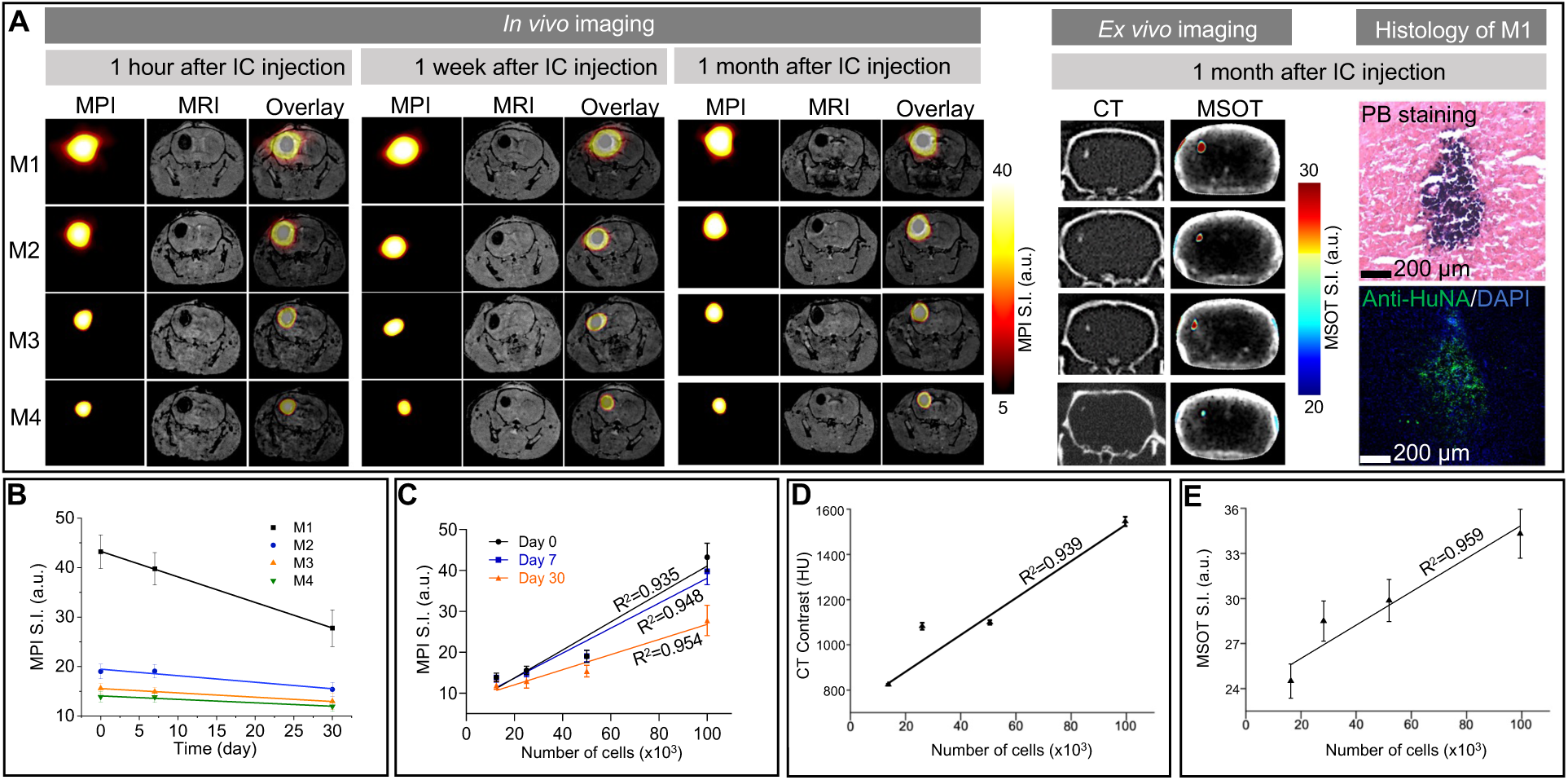
Serial *in vivo* and *ex vivo* imaging and quantitative analysis of ABS-labeled hMSCs. (**A**) *In vivo* MPI/MRI of mice injected with varying numbers of ABS-labeled hMSCs at different time points post-injection. M1, M2, M3, and M4 received 1.00×10^5^, 5.00×10^4^, 2.50×10^4^, and 1.25×10^4^ cells, respectively. *Ex vivo* CT and MSOT imaging, along with histology of M1 (as a representative example), were also performed to further validate ABS-labeled hMSC distribution and localization. (**B**) Time-course analysis of *in vivo* MPI signal intensity, showing a signal decay over 30 days for different cell quantities. (**C**-**E**) Correlation between *in vivo* MPI, *ex vivo* CT, and *ex vivo* MSOT signal intensity and the number of injected cells, demonstrating strong linear relationships.

A time-course analysis of MPI signal intensity demonstrated a signal decay over 30 days (**Fig. 5B**). This decay may either result from cell death or the exocytosis of ABS from hMSCs, with subsequent clearance and breakdown by macrophages^4^. As our study primarily focused on evaluating the potential of ABS for stem cell tracking using clinically relevant imaging techniques, we did not include bioluminescence imaging (BLI). Clinical MRI, CT, and MSOT machines are available, while MPI is now in clinical development for human head-sized scanners^39–44^ and handheld devices^45^.

**Fig. 5C-E** present the quantitative correlation between the number of ABS-hMSCs injected and the signal intensity recorded for MPI, CT, and MSOT respectively, showing the capacity of each imaging modality to quantitatively assess cell number. This strong linear relationship allows accurate quantitative stem cell tracking.

## Conclusions

We introduced ABS as a novel nanocomplex for multimodal anatomical and quantitative imaging of stem cells, leveraging the unique properties of albumin, Bi_2_S_3_ NPs, and ferucarbotran. Through comprehensive physicochemical characterization, we established the robust stability of ABS in terms of structural integrity and functional performance. Key findings include the preservation of structural components of albumin within the ABS nanocomplex and the incorporation of SPIO and Bi_2_S_3_ without altering the primary BSA protein structure. We found that CT, MPI, and MSOT have distinct sensitivity thresholds for quantifying both naked ABS and ABS-labeled cells. MPI demonstrated the greatest sensitivity, followed by MSOT, and then CT, while MRI was found not to be quantitative Our results highlight the ABS capability to address some of the existing limitations of MRI, CT, MPI, and MSOT. The integration of MPI and MRI provided a dual approach to visualize cells in a quantitative manner within their anatomical context, a significant advancement over single-modality magnetic imaging. With interventional radiology cell-injection procedures most commonly performed using CT or fluoroscopy, the inclusion of Bi_2_S_3_ in ABS not only enhances the sensitivity of CT but also enables sensitive real-time stem cell tracking using MSOT.

Looking forward, ABS may present numerous opportunities to develop new imaging-guided precision cell therapies. Integrating ABS with advanced technologies such as targeted drug delivery systems and real-time monitoring of therapeutic interventions could significantly advance cell therapy.

## Methods

### Materials for synthesis of ABS

Ferucarbotran (stock concentration 1 M Fe) was purchased from Meito Sangyo.

BSA (Cat. # A9647, Lot # SLCH8436, pH 7, ≥98% purity), bismuth (III) nitrate pentahydrate (Bi(NO_3_)_3_.5H_2_O), 70% nitric acid, and NaOH were purchased from Sigma-Aldrich, and 1-ethyl-3-(3ʹ-dimethylaminopropyl)carbodiimide, hydrochloride (EDAC, HCl) or C_8_H_17_N_3_ · xHCl was purchased from EMD Millipore Corp. Lot variations of BSA were observed to result in slight size variations of AB and ABS, but all data were acquired with high consistency using the specific BSA lot listed above.

### Synthesis of BSA-Bi_2_S_3_ (AB) nanocomplexes

Three solutions were prepared. **Solution 1** was prepared by dissolving 13.2 g of BSA in 200 ml of double-distilled water (ddH₂O) in a glass container under continuous stirring until all BSA granules were completely dissolved. **Solution 2** was prepared by dissolving 17 mg of Bi(NO₃)₃·5H₂O in 2.88 ml of 70% HNO₃ in a 15 ml tube. After the Bi salt was completely dissolved, 11.12 ml of ddH₂O was added, and the mixture was shaken to achieve a homogeneous solution. **Solution 3** was prepared by dissolving 6 g of NaOH in 30 ml of ddH₂O in a 50 ml tube at pH=12.75). While **Solution 1** was being stirred vigorously in the glass container, **Solution 2** was added dropwise. If white precipitation appeared during this process, the solution was homogenized using an ultrasonic bath to ensure a fully transparent yellow solution. Finally, **Solution 3** was added to the mixture of Solutions 1 and 2 while stirring vigorously. After 30 mins of stirring, the color of the mixture changed from light yellow to black. The mixture was then stirred gently for an additional 8 h. The final product was collected by centrifuging the mixture at 13,500xg for 10 min, followed by washing the precipitate three times with ddH₂O. The yield was approximately 87%, based on bismuth content.

### Synthesis of BSA-Bi_2_S_3_-SPIO (ABS) nanocomplexes

Six mg of EDAC was mixed with 2 ml of SPIO (Ferucarbotran, 1 mg Fe/ml), followed by the addition of 12 ml of ddH₂O to bring the total volume to 14 ml. The mixture was shaken for 15 min. Subsequently, 2 ml of AB (5 mg Bi/ml) was added, and the mixture was shaken for an additional 2 h. The final product was washed three times with ddH₂O using centrifugation at 6,400xg for 5 min, employing a centrifugal filter tube (Amicon Ultra-15, 10kDa MWCO) for purification. The collected ABS nanocomplexes were then sonicated for 15 min. The yield was approximately 90%, based on Fe content.

### Physicochemical characterization of ABS

A Shimadzu UV-1900i spectrophotometer and a Shimadzu FT-IR 4300 instrument were used for optical characterization. A Malvern Zetasizer Nano ZS instrument with back scattering detector (173°) was used for measuring the hydrodynamic size and zeta potential in batch mode. The following NIST-NCL joint protocol PCC-1 was followed: https://www.cancer.gov/nano/research/ncl/protocols-capabilities. Stock samples were diluted 100-fold in water, 10 mM NaCl, and PBS. Samples were measured at 25 °C in a quartz microcuvette. Zeta potential measurements were made at native pH and, where necessary, after adjustment to near neutral pH (7.4) using either 1 N standardized HCl or 1 N standardized NaOH. Sample pH was measured and/or adjusted before loading into a pre-rinsed folded capillary cell.

Particle size distribution was also measured using AF4 coupled with MALS and DLS detectors. The AF4 system consisted of an isocratic pump (Agilent G1310A), well-plate autosampler (Agilent G1329A), UV-vis detector (Agilent G1315B), AF4 separation channel (DualTec, Wyatt Technology), MALS detector (Wyatt HELEOS II), refractive index (RI) detector (Wyatt OptiLab T-rEX), and a DLS (Malvern Zetasizer Nano ZS) instrument. The separation channel had a length of 275 mm and a 350 µm spacer. A 10kDa MWCO regenerated cellulose membrane was used for particle separation. The detector flow was 1 ml/min and the injection volume was 100 µl for all samples. The mobile phase was PBS. The elution cross-flow was 1.50 ml/min for 10 min, followed by a linear decrease to 0 ml/min in 3 min, and held at 0 ml/min for 25 min. The membrane was passivated with 5 mg/ml BSA for a total of three runs prior to sample analysis. The chromatographic traces were monitored by absorption at 210 nm, MALS, and DLS detection. MALS normalization constants were determined using BSA at 5 mg/ml in PBS. DLS was used for measuring the hydrodynamic diameter in flow-mode. Samples were diluted 100-fold in PBS prior to injection. For plasma incubation experiments, 5 µl of sample were incubated with 100 µl of human plasma at 37 °C for 2 h with agitation. Then, 895 µl PBS was added to the mixture, which was further diluted 1:1 with PBS for a final 400-fold dilution in 5% human plasma.

The hydrodynamic size and particle concentration were also measured using a Wyatt DynaPro plate reader III (Wyatt Technology). The instrument was calibrated with dextran standards and the PBS solvent was filtered through a 0.02 μm regenerated cellulose membrane prior to use. The size and particles per ml concentration were also measured using NTA (ViewSizer 3000). Measurements utilized all laser lines. Samples were diluted in water accordingly (100,000 to 2,000,000-fold) to give a particle count of approximately 107 particles/ml.

The iron, sulfur and bismuth concentrations in ABS and AB were determined by ICP-MS. A Perkin-Elmer NexION 2000B equipped with a micro-mist nebulizer, standard sample introduction system, and integrated auto-sampler operated in standard mode was used. Data was analyzed using the Syngistics software. Samples for Bi and Fe analysis were prepared by serial dilution with three dilution steps. Dilution 1:50 µl of sample was weighed out and digested in 400 µl HNO_3_ and 100 µl of HCl, followed by dilution to 10 ml using a 2% HNO_3_ solution. Dilution 2:100 µl of Dilution 1 was weighed out and diluted to 10 ml using 2% HNO_3_. Dilution 3:1 ml of Dilution 2 was weighed and diluted to 10 ml using 2% HNO_3_. Total weights of the diluted solutions were also recorded and used to determine the dilution factors and calculate the amount of Fe or Bi in solution. Samples for sulfur analysis were prepared by single dilution. Dilution 1:50 µl of sample was weighed out and diluted in 10 ml of Milli-Q Water. Iron calibration standards were prepared from a serial dilution of a 0.997 mg/g Fe in HNO_3_ (Inorganic Ventures). The iron calibration standards prepared had concentrations of 0, 2.186, 4.367, 6.845, 8.668, and 11.10 ng/g. Bismuth calibration standards were prepared from a serial dilution of a 0.997 mg/g Bi in HNO_3_ (Inorganic Ventures). The bismuth calibration standards prepared had concentrations of 0, 10.40, 20.65, 30.85, 41.28, and 51.81 ng/g. Sulfur calibration standards were prepared from a serial dilution of a 0.997 mg/g S (Inorganic Ventures) in water. The sulfur calibration standards prepared had a concentration of 0, 0.9688, 5.001, 10.13, 30.13, and 50.28 ng/g.

All concentrations were determined using an external calibration constructed using the standards above. Iron, bismuth, and sulfur were measured using peak hopping with an integration time of 50 ms. Each measurement consisted of 3 sweeps with 6 reading per sweep, and the final Intensity was determined from an average of 10 replicants for each sample. Samples were run in KED mode with He as the reaction gas. Masses analyzed included Bi-209, Fe-57, and S-34.

### Labeling of hMSCs

P2 human bone marrow-derived MSCs were obtained from RoosterBio™ and expanded in culture up to P5 using MSC basal medium (MSCBM, Lonza) supplemented with 10% MSC growth supplement, 2% l-glutamine, 0.1% gentamicin, and 0.1% amphotericin. For all labeling experiments except the cell viability assay, hMSCs were incubated for 24 h with a pre-prepared labeling medium consisting of MSCBM medium supplemented with either ABS (25 µg Fe/ml), AB (125 µg Bi/ml), or ferucarbotran (25 µg Fe/ml), along with poly-L-lysine (Sigma P-1524) as a cationic agent at a concentration of 3125 ng/ml. Following incubation, the labeling medium was removed, and the labeled hMSCs were washed three times with PBS to remove unbound particles. TrypLE™ Express (Gibco) was used to gently detach the labeled cells for subsequent staining or injection into mice.

### Prussian blue staining

PB staining was performed to detect the presence of iron within labeled hMSCs. Cells were fixed with 4% glutaraldehyde for 20 mins, washed with PBS, and incubated with Perl’s reagent (potassium ferrocyanide in HCl) for 30 min at room temperature. To enhance cellular visualization, cells were counterstained with nuclear fast red for 20 mins. Stained cells were washed with deionized water to remove excess reagent and then imaged using a Zeiss Apotome 2 microscope.

### Ferrozine assay

The Fe concentration in naked NPs or labeled hMSC was determined using a Ferrozine assay as described elsewhere^46^. A standard curve prepared with known Fe concentrations was used to calculate the Fe content in the sample.

### Cell viability assay

To evaluate the potential cytotoxicity of ferucarbotran, AB, and ABS, a lactate dehydrogenase (LDH) assay (ThermoFisher Scientific) was performed for 24 or 72 hr incubation with various NP concentrations. hMSCs were seeded in 96-well plates and following incubation the culture supernatant was collected, and the absorbance was measured at 490 nm according to the manufacturer’s protocol. The cytotoxicity was expressed as percentage of the absorbance relative to the positive control (100% cell death).

### TEM

Nanocomplexes (10 μl) were adsorbed on glow-discharged ultra-thin carbon-coated 400 mesh copper grids (EMS CF400-Cu-UL). Cells were fixed in 4% paraformaldehyde (PFA), 0.1% glutaraldehyde, 3 mM MgCl_2_ in and 0.1 M Sorenson’s sodium phosphate buffer, pH=7.2 overnight at 4 °C. After buffer rinse, samples were postfixed in 1% osmium tetroxide and 1.5% potassium ferrocyanide in 0.1 M sodium phosphate for 1h on ice in the dark. Samples were rinsed in ddH_2_O, dehydrated in a graded series of ethanol and embedded in Eponate 112 (Polyscience) resin, and polymerized at 60 °C overnight. Sixty to 90 nm sections were cut with a diamond knife on a Leica UCT ultramicrotome and collected with Formvar coated 2×1 copper slot grids. Before imaging, grids were stained with 2% uranyl acetate, rinsed with ddH_2_O and stained with 0.3% Reynolds lead citrate. Grids were imaged with a Hitachi 7600 TEM at 80 kV equipped with an AMT CCD XR80 (8-megapixel camera - side mount AMT XR80 - high-resolution high-speed camera).

HAADF-STEM imaging was performed using a JEOL JEM-ARM200F microscope operated at 200 kV to visualize the morphology and structural features of ABS. For elemental analysis, EDS mapping for Fe, Bi and S was conducted using an Oxford Instruments X-MaxN detector integrated into the microscope.

### Labeled cell injections

All animal experiments were approved by our Institutional Animal Care and Use Committee. Male immunodeficient male Rag2^−/−^ mice (6–8 weeks old, The Jackson Laboratory) were housed under a 12-h light/dark cycle with unrestricted access to food and water. Intracerebral injections were performed under 1%–2% isoflurane anesthesia with mice secured in a stereotaxic frame (Stoelting Co.). A burr hole was created 0 mm caudal and 2 mm lateral to the bregma. Using a 31G Hamilton syringe (Hamilton), 2 μl of cell suspension in PBS was injected into the striatum at a depth of 3 mm below the endocranium. Varying quantities of cells (1×10^5^, 5×10^4^, 2.5×10^4^, and 1.25×10^4^ cells) were delivered at a rate of 0.2 μl/min over 10 min, with the needle left in place for 5 mins before withdrawal. The incision was closed using 3-0 vicryl sutures, and 1 mg/kg buprenorphine ER was administered s.c. once post-surgery.

### Quad-modal imaging

To directly compare the *in vitro* sensitivity of the four imaging modalities, naked AB, naked ABS and ABS-labeled hMSC sample were prepared at different cell densities. Naked NP samples were suspended in 100 µl of DI water in small 200 µl Eppendorf tubes. Labeled cell samples were prepared in 100 µl of 8% gelatin with 0.02% w/v sodium azide added. *In vivo* MRI and MPI were performed at 1 hr, 1 week, and 1 month after ABS-hMSC injection. Following *in vivo* MRI/MPI imaging, mice were sacrificed and *ex vivo* MSOT/CT imaging were conducted, followed by post-mortem analysis.

### MRI

MRI was conducted using a Bruker 11.7 T Bruker Biospin horizontal bore scanner equipped with a 25-mm-volume coil and interfaced with ParaVision 6.0.1 software. *In vitro* MRI was performed to generate 2D T2-w images using a multi-slice multi-echo sequence. *In vitro* MRI parameters included an echo spacing of 6.5 ms, a repetition time (TR) of 2200 ms, and a slice thickness of 1.5 mm. The image matrix size was 192×192, with a field of view (FOV) of 40 ×40 mm, resulting in an in-plane resolution of 0.2×0.2 mm. Each *in vitro* scan used one average and one repetition.

A 3D FLASH sequence was employed for *in vivo* MRI studies, featuring isotropic imaging with a slice thickness of 40 mm and an image matrix size of 200×125×250. The FOV was 32×20×40 mm, with a resolution of 0.16×0.16×0.16 mm. Scan parameters were a TE=3.4 ms, a TR=25 ms, and a flip angle of 7° with fat suppression. Each *in vivo* scan used a single average with one repetition. Regions of interest (ROIs) were manually drawn and data were processed using ImageJ.

### MPI

MPI was performed using a Momentum field-free line imager (Magnetic Insight Inc.). Phantoms were positioned vertically within a 3D custom-printed holder (Ultimaker 2 Extended+), and 2D images were acquired using the manufacturer’s “standard” settings. For samples with lower iron concentrations or cell densities, tubes were imaged individually by placing eppendorf tubes vertically in the center of the holder, aligned with the MPI FOV.

For *in vivo* imaging, a 2D whole-body MPI scan was initially performed in standard mode, followed by 3D MPI scan (standard mode) with 21 projections ROIs were manually drawn and data were processed using ImageJ.

### MPI/MRI data co-registration

Co-registration was performed using the Volume, Look-Up Table, Transform, and Rendering tools in 3D Slicer. To facilitate co-registration and quantification, two fiducial tubes containing ABS-labeled hMSCs, prepared from the same batch of injected cells, were included in the imaging setup. These fiducials were then used as reference points for accurate alignment of MPI and MRI datasets.

### MSOT

MSOT imaging was performed using an MSOT inVision 512-echo scanner (iThera Medical), equipped with a 512- toroidally-focused ultrasound transducer array operating at a central frequency of 5 MHz, spanning a circular arc of 270° to effectively detect optoacoustic signals. Light excitation was provided with a tunable optical parametric oscillator pumped by an Nd:YAG laser. Excitation pulses with a duration of 9 ns at wavelengths ranging from 660 to 900 nm at a repetition rate of 10 Hz, wavelength tuning speed of 10 ms, and a peak pulse energy of 100 mJ at 720 nm were used.

Sample tubes were embedded in cylindrical phantoms^47,48^.

For *ex vivo* MSOT, mice were first transcardially perfused with 10 mM PBS and 4% PFA. Heads were collected, shaved, and wrapped in a thin polyethylene membrane and then placed in a customized animal holder (iThera Medical). A thin layer of ultrasound gel was applied to the skin for optimal acoustic coupling to the membrane prior to imaging. MSOT data were acquired with excitation wavelengths ranging from 660 to 900 nm in steps of 5 nm, with 10 frames recorded and averaged for each wavelength. All MSOT data analysis was performed using viewMSOT software (Version 4.0.2.0). A linear regression method was used to perform multispectral processing.

MSOT images were reconstructed from the raw data using a back-projection algorithm at a resolution of 100 μm. ROIs were manually drawn based on concurrently acquired b- Mode ultrasound images. Within each ROI, the mean of the highest 10% of pixels was used for quantification. Analyses were computed with a NP unmixing preset, using the spectrum derived from a sample with highest NP concentration.

### Micro-CT

An IVIS Spectrum/CT imaging system (Caliper Sciences) was used with a tube voltage of 50 kVp, a current of 200 µA, and an exposure time of 300 ms. Images were acquired at a voxel size of 100 µm, and 3D reconstructions were generated using the accompanying software. Images were processed using ImageJ. To quantify CT signal intensity as HU, a two-point calibration method was used. The CT signal intensity in water was set to 0 HU, and that for ambient air to −1000 HU. Sample HU values were then obtained by linear extrapolation using

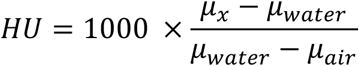

where, μ_x_, μ_water_, and μ_air_ are the linear attenuation coefficients of the sample, water, and air, respectively.

### Histopathology

After *ex vivo* imaging, the brains were removed and immersed into 4% PFA at 4 °C for 24 h, and then transferred to 30% sucrose for another 72 h. After embedding in optimal cutting temperature compound, the brains were sequentially cut into 20 μm sections using a cryostat (Thermo Fisher Scientific). For PB staining, tissues were fixed with 4% glutaraldehyde for 20 min and staining was performed as described above.

Anti-human nuclear antigen (anti-HuNA) staining was performed to visualize hMSCs. Tissue sections were first blocked with 5% BSA and 0.1% Triton X-100 in Tris-buffered saline (×TBS, pH=7.0) for 2 h, followed by incubation with mouse anti-HuNA (1:250, NBP2-34342, Novus-Bio). Goat anti-mouse Alexa-fluor 488 (1:500, A-11001, Invitrogen) was used as secondary antibody, prepared in 3% BSA in TBS. Sections were incubated for 2 h at room temperature and then triple washed in 1×TBS to remove antibody for each step. Sections were cover-slipped with mounting medium containing DAPI. Fluorescence microscopy was performed using a Zeiss Axiovert 200 M inverted epifluorescence microscope.

## Statistical analysis

Linear regression analysis was performed to plot the total signal (a.u.) from each imaging modality against NP concentration or labeled cell density. The R² value of each regression line was used to evaluate the linearity between the signal and either NP concentration or cell density. Data are expressed as the mean ± standard deviation from at least three independent experiments (technical replicates). Statistical analysis was conducted using GraphPad Prism software (version 10.3.0). Comparisons across multiple groups were carried out using one-way analysis of variance followed by Tukey’s post hoc test, with a significance level of p<0.05.

## Data Availability

Data are available from the authors upon request.

## Acknowledgements

This work was funded by grants from the National Institutes of Health (R01 CA257557, UH2/UH3 EB028904 and S10 OD026740) and the Maryland Stem Cell Research Foundation (MSCRFD-5416, MSCRFL-6270). This work was also supported, in part, by federal funds from the National Cancer Institute, National Institutes of Health, under contract no. 75N91019D00024. The formulation described herein was accepted into the Assay Cascade characterization program of the Nanotechnology Characterization Laboratory (NCL) of the Frederick National Laboratory for Cancer Research. The NCL provides a free characterization service for cancer-related nanomedicine formulations, available to the public by application (https://www.cancer.gov/nano/research/ncl). The content of this publication does not necessarily reflect the views or policies of the Department of Health and Human Services, nor does mention of trade names, commercial products, or organizations imply endorsement by the U.S. Government.

