## Supplemental Info for "Cold Quad-Modal Nanocomplex for Precise and Quantitative *in Vivo* Stem Cell Tracking"

**Particle sizing and concentration**

The hydrodynamic size and distribution were measured for AB and ABS after dilution in water only (without salt),10 mM NaCl in water, or 10 mM phosphate buffered saline, pH=7.4 (PBS). The intensity-weighted peak for ABS was approximately 136 nm when diluted in all three-dispersing solutions (**Fig. S1, Table S1**). The intensity-weighted peak for the precursor formulation, AB, was approximately 105 nm when diluted in 10 mM NaCl and PBS (**Fig. S2, Table S2**). When diluted in water only, the size increased to approximately 122 nm. The AB particles were smaller than ABS, as expected due to the addition of the SPIO (ferucarbotran) to the surface of AB. Ferucarbotran had an intensity-weighted peak of approximately 85 nm when diluted in 10 mM NaCl and PBS. The size increased to approximately 120 nm when diluted in water only (**Fig. S3, Table S3**).

The size and particle concentration were measured for ABS, AB, and ferucarbotran using a Wyatt DynaPro plate reader, based on light scattering. ABS had an average size of 120.0 nm and a concentration of 2.9x$10^{12}$ particles/mL (**Fig. S4A, Table S4**). AB had an average size of 87.1 nm and a concentration of 6.4x$10^{12}$ particles/mL (**Fig. S4B, Table S5**). Ferucarbotran had an average size of 67.5 nm and a concentration of 2.35x$10^{12}$ particles/mL (**Fig. S4C, Table S6**). The particle concentration was also measured for ABS using a Horiba ViewSizer 3000, based on NTA. ABS had an average mean and mode size of 99.1 and 76.6 nm, respectively, with a concentration of 2.47x$10^{13}$ particles/mL (**Fig. S5, Table S7**).

**Quantitative elemental analysis**

The total Bi and Fe concentration in ABS and AB were measured using ICP-MS as listed in **Table S8**. ABS had a Bi concentration of 5.09±0.52 and an Fe concentration of 1.23±0.05 mg/mL. Sulfur concentrations in ABS, AB and BSA were measured using ICP-MS. The sulfur content was then used to determine the amount of BSA in AB and ABS. AB and ABS had sulfur concentrations of 1.35±0.08 and 1.73±0.08 mg/mL, respectively (**Table S8**). Assuming 32 sulfur atoms per BSA molecule, the measured sulfur concentrations were converted to equivalent BSA concentrations. ABS and AB had BSA concentrations of 84.6±5.0 and 108.5±5.0 mg/mL, respectively. The S:Bi ratio was 1.14 and 1.33 for ABS and AB, respectively. Single particle ICP-MS was used to determine the particle concentration in ABS and AB (**Table S9**). ABS and AB had particle concentrations of 1.72x$10^{13}$ and 1.97x$10^{13}$ particles/mL, respectively (**Table S10**).

**Zeta potential analysis**

ABS and AB had negative zeta potential values at native pH (**Fig. S6, Table S11**). Ferucarbotran was neutral at its native pH, as zeta potentials ranging from +10 to -10 mV are generally considered neutral. The negative zeta potential for ABS suggests low ferucarbotran loading per particle and that the dextran coating is possibly interweaved within the BSA coating.

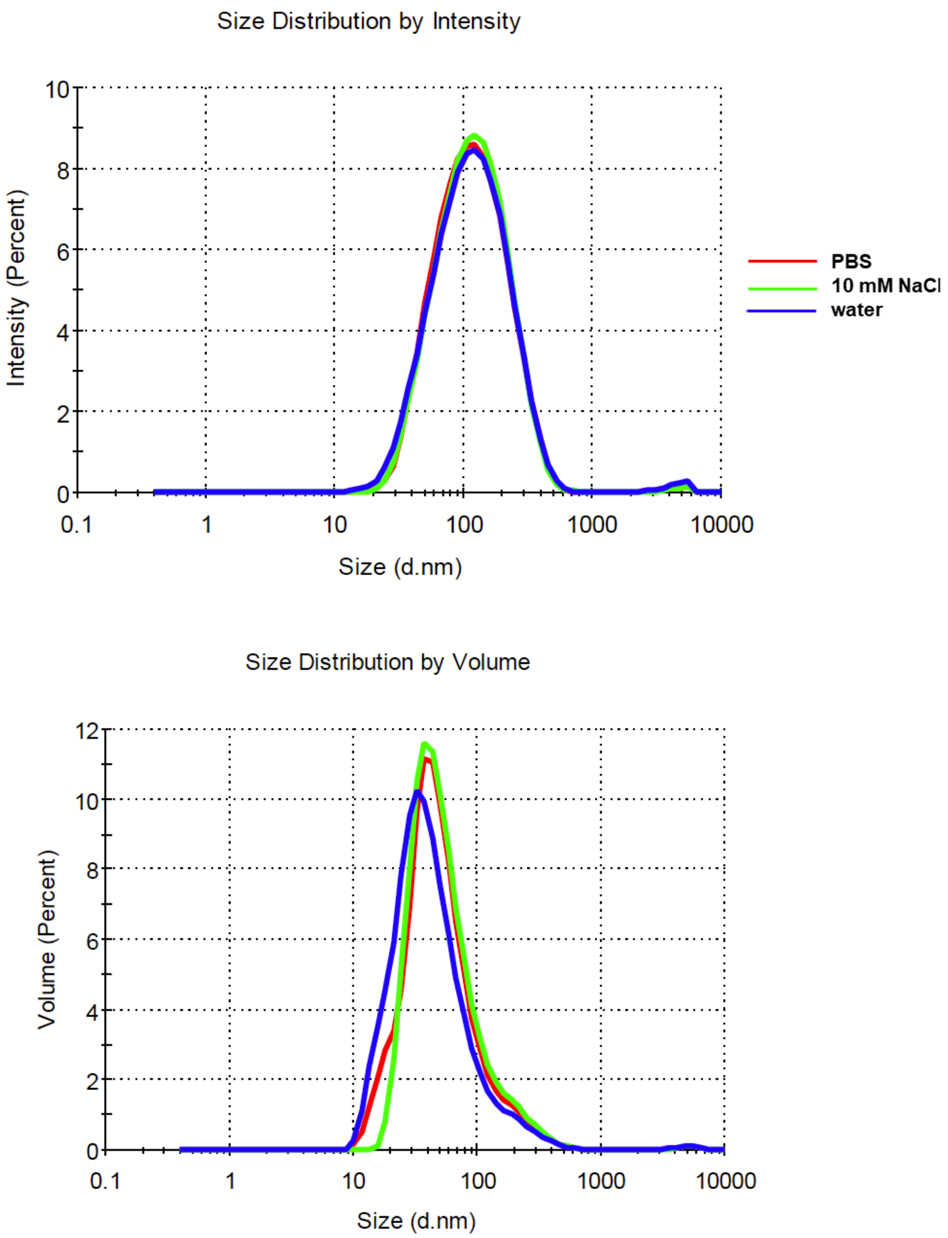


**<u>Figure S1:</u>** Representative DLS analysis showing the averaged intensity and volume distribution plots for ABS.

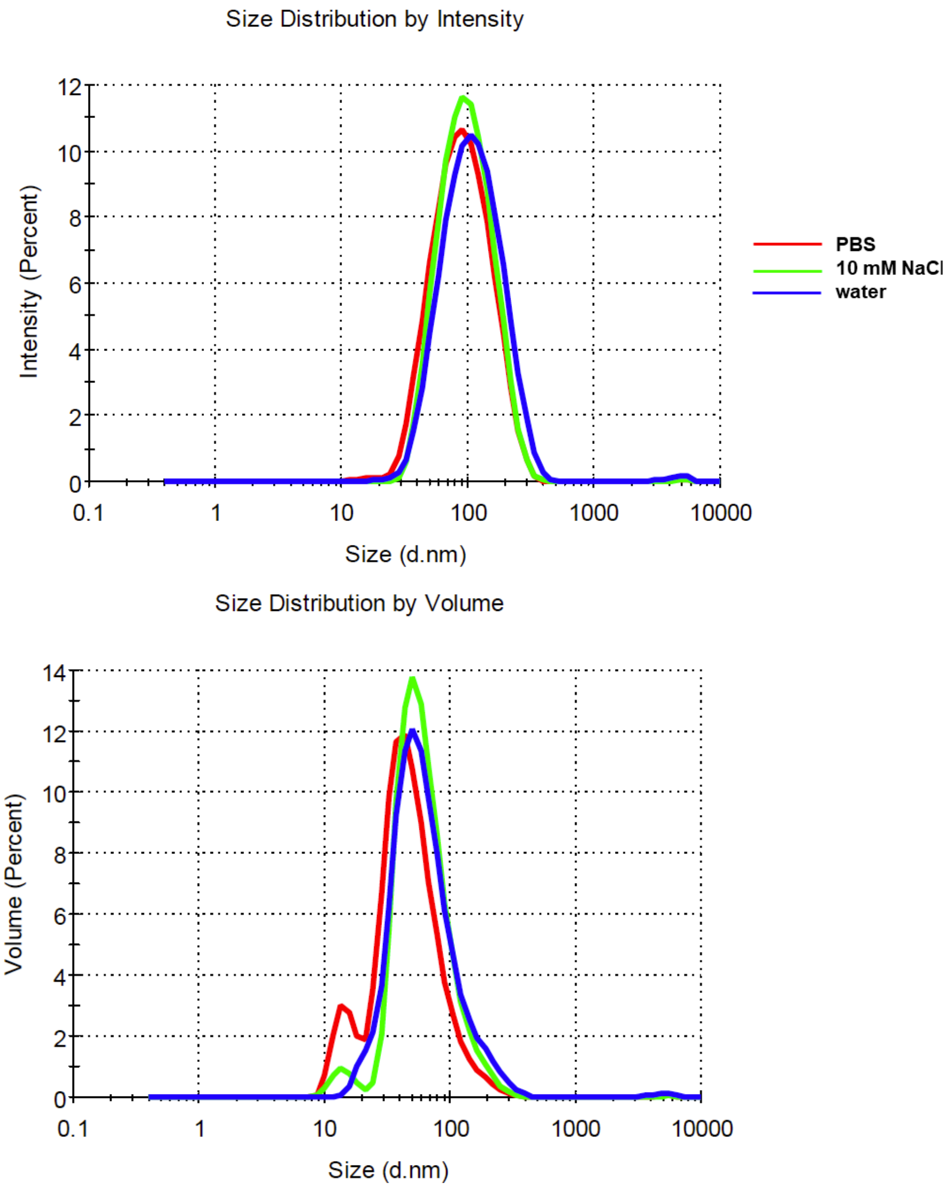


**Figure S2:** Representative DLS analysis showing the averaged intensity and volume distribution plots for AB.

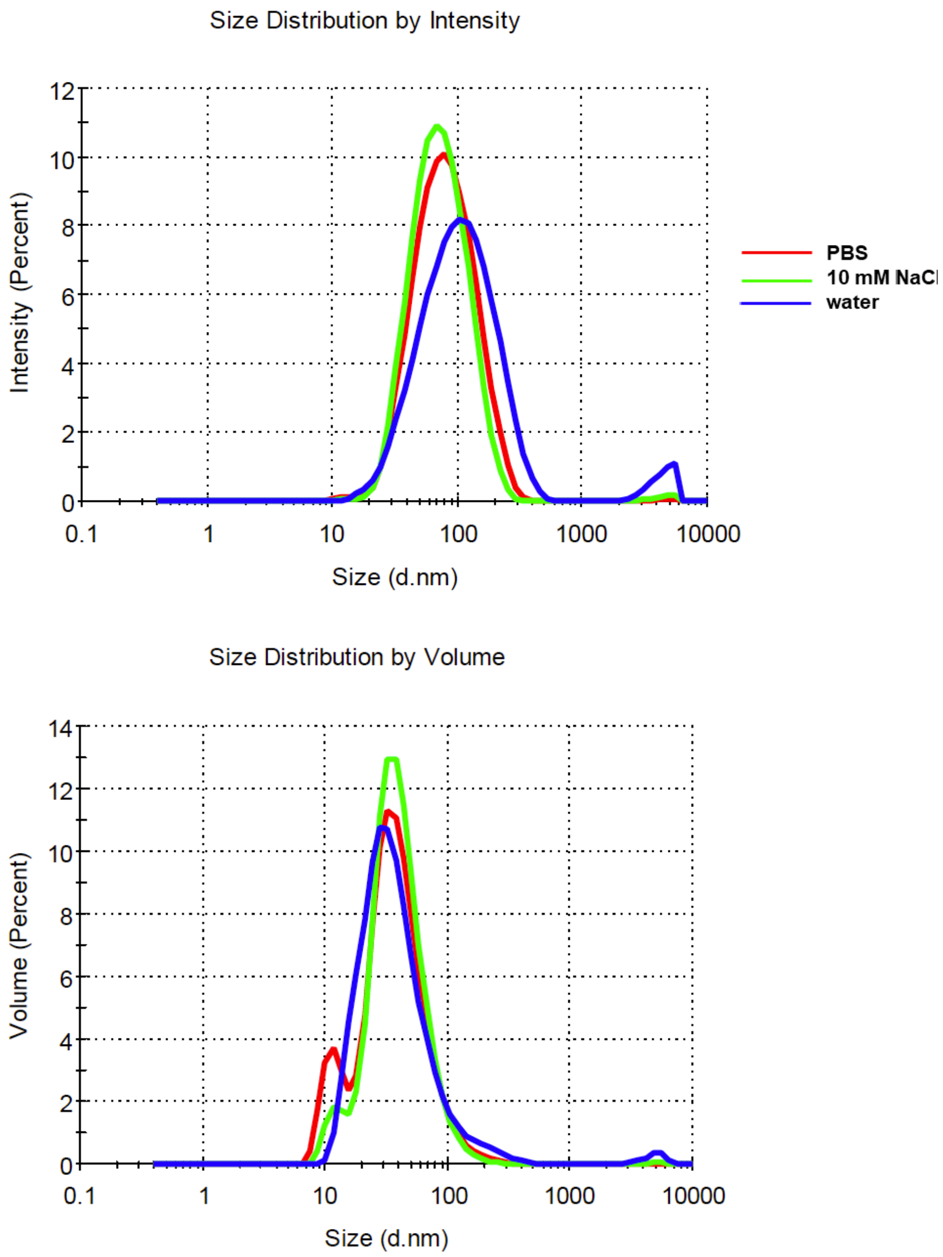


**<u>Figure S3:</u>** Representative DLS analysis showing the averaged intensity and volume distribution plots for ferucarbotran.

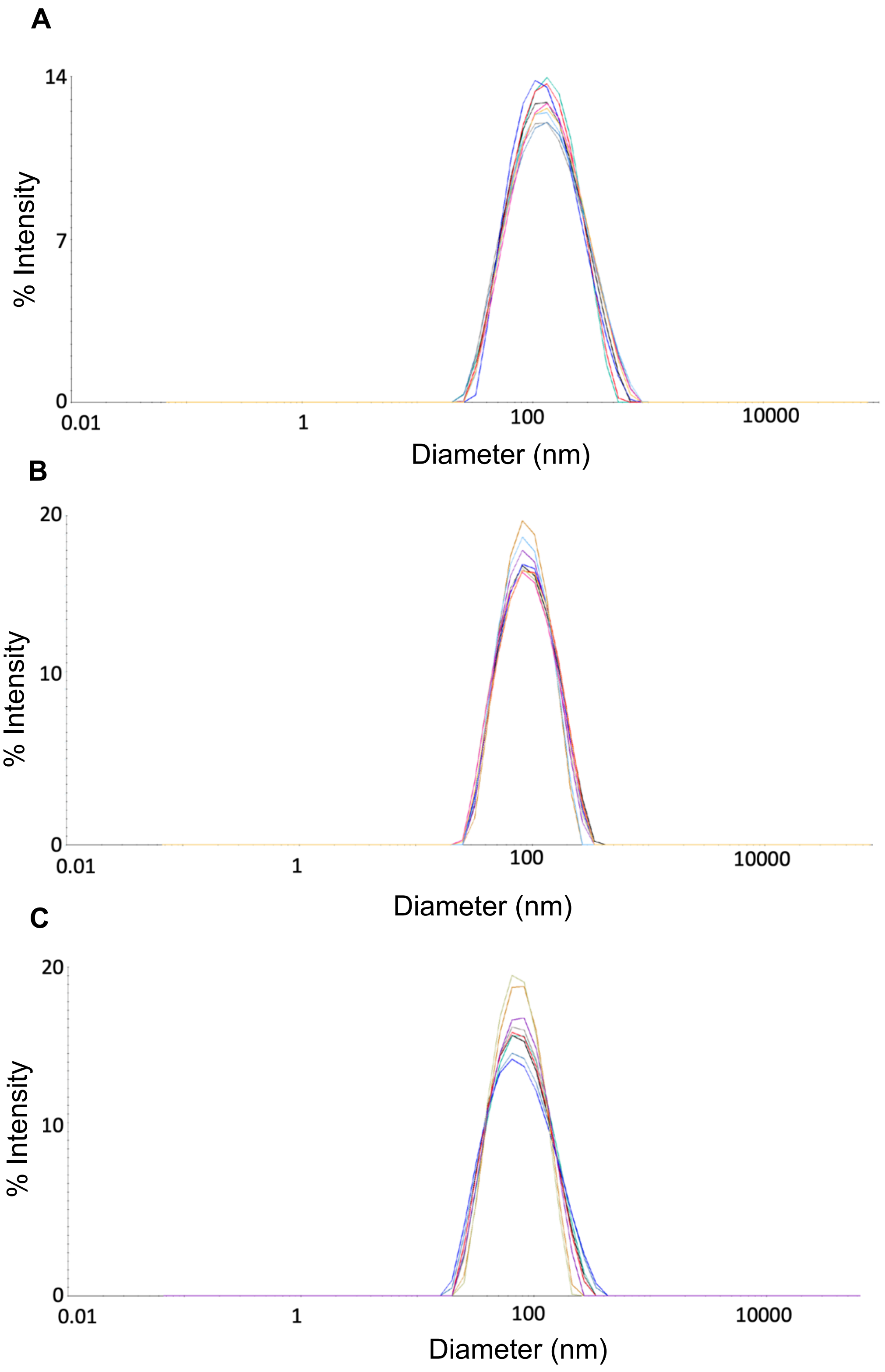


**<u>Figure S4:</u>** Replicate hydrodynamic size distributions for (**A**) ABS, (**B**) AB, and (**C**) ferucarbotran, as determined by using a Wyatt DynaPro plate reader.

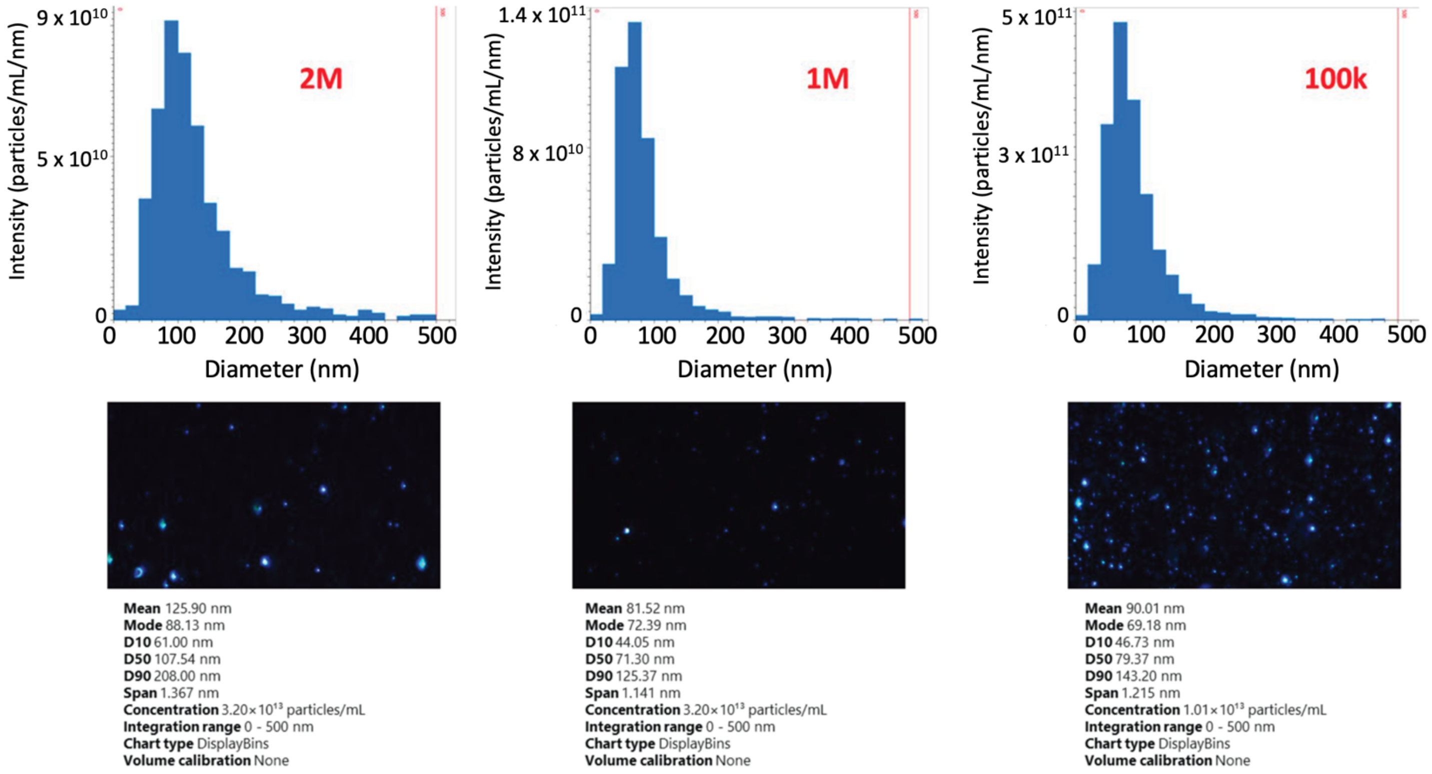


**Figure S5:** Size distribution and representative sample image of ABS for three dilutions using NTA. Samples were diluted 2,000,000 (2M), 1,000,000 (1M) and 100,000 (100k) fold in water before analysis.

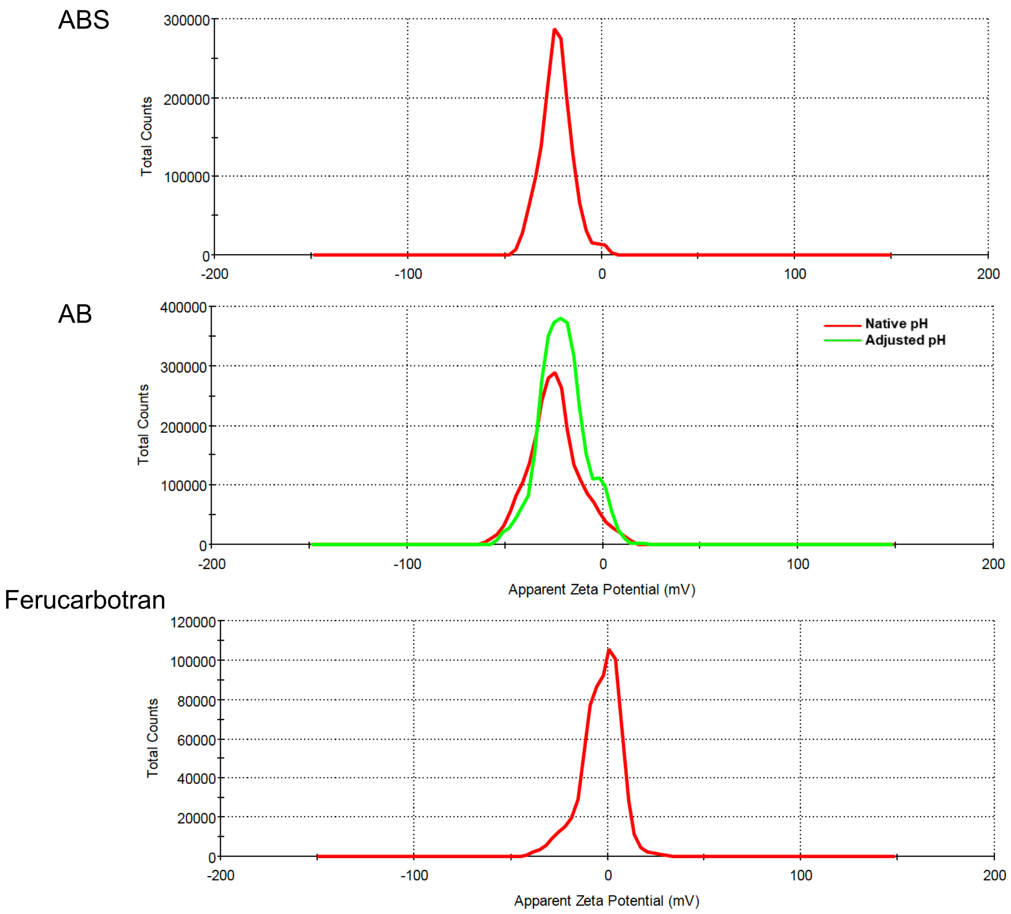


**Figure S6:** Averaged zeta potential distributions for ABS (pH 7.29), AB (pH 9.25 native, pH 7.59 adjusted), and ferucarbotran (pH 7.42 native).

**Table S1:** Hydrodynamic diameter of ABS. PdI = Polydispersity index, Int = Intensity.

| Sample | Dilution | Z-Avg (nm) | PdI | Intensity-based (nm) | % Int | Volume-based (nm) | % Vol |
|---|---|---|---|---|---|---|---|
| ABS | 1:100 in PBS | 97.4 ± 1.2 | 0.267 ± 0.006 | 136.1 ± 5.8 | 99.3 ± 1.1 | 64.0 ± 8.0 | 99.6 ± 0.7 |
| ABS | 1:100 in 10 mM NaCl | 99.0 ± 1.0 | 0.263 ±0.005 | 137.5 ± 7.2 | 99.6 ± 0.9 | 66.2 ± 8.9 | 99.7 ± 0.7 |
| ABS | 1:100 in water | 94.5 ± 0.8 | 0.278 ± 0.006 | 136.2 ± 7.2 | 99.1 ± 1.0 | 56.3 ± 7.9 | 94.0 ± 13.5 |

**Table S2:** Hydrodynamic diameter of AB. PdI = Poly dispersity index, Int = Intensity.

| Sample | Dilution | Z-Avg (nm) | PdI | Intensity-based (nm) | % Int | Volume-based (nm) | % Vol |
|---|---|---|---|---|---|---|---|
| AB | 1:100 in PBS | 79.4 ± 0.3 | 0.211 ± 0.0047 | 102.1 ± 2.7 | 99.5 ± 0.8 | 56.7 ± 3.1 | 88.8 ± 20.7 |
| AB | 1:100 in 10 mM NaCl | 86.3 ± 1.3 | 0.200 ± 0.010 | 107.4 ± 2.9 | 99.8 ± 0.4 | 67.9 ± 3.2 | 96.4 ± 12.1 |
| AB | 1:100 in water | 95.5 ± 3.2 | 0.223 ± 0.008 | 122.2 ± 6.4 | 99.4 ± 1.1 | 72.3 ± 6.6 | 95.9 ± 12.8 |

**Table S3:** Hydrodynamic diameter of ferucarbotran. PdI = Poly dispersity index, Int = Intensity.

| Sample | Dilution | Z-Avg (nm) | PdI | Intensity-based (nm) | % Int | Volume-based (nm) | % Vol |
|---|---|---|---|---|---|---|---|
| Ferucarbotran | 1:100 in PBS | 68.4 ± 1.3 | 0.235 ± 0.009 | 90.6 ± 4.0 | 99.5 ± 0.8 | 45.3 ± 4.8 | 87.3 ± 23.9 |
| Ferucarbotran | 1:100 in 10 mM NaCl | 63.5 ± 0.8 | 0.226 ± 0.009 | 80.6 ± 3.2 | 99.3 ± 1.2 | 44.5 ± 3.4 | 92.8 ± 16.9 |
| Ferucarbotran | 1:100 in water | 86.9 ± 3.1 | 0.368 ± 0.014 | 120.2 ± 7.7 | 95.3 ± 2.9 | 51.5 ± 10.3 | 86.5 ± 28.8 |

**Table S4:** Hydrodynamic diameter and nanoparticle concentration for ABS. Values were corrected for dilution.

| Sample | Diameter (nm) | Attenuation Level (%) | Laser Power (%) | Particle Concentration (particles/mL) |
|---|---|---|---|---|
| ABS | 117.9 | 90 | 60 | $3.50\times10^{12}$ |
| ABS | 120.4 | 90 | 60 | $2.82\times10^{12}$ |
| ABS | 121.7 | 90 | 80 | $2.39\times10^{12}$ |
| **Average** | **120.0** | | | **$2.90\times10^{12}$** |
| **SD** | **1.6** | | | **$4.59\times10^{11}$** |

**Table S5:** Hydrodynamic diameter and nanoparticle concentration for AB. Values were corrected for dilution.

| Sample | Diameter (nm) | Attenuation Level (%) | Laser Power (%) | Particle Concentration (particles/mL) |
|---|---|---|---|---|
| AB | 88.0 | 75 | 100 | $5.11x10^{12}$ |
| AB | 86.6 | 75 | 60 | $9.06x10^{12}$ |
| AB | 86.6 | 50 | 60 | $5.02x10^{12}$ |
| **Average** | **87.1** | | | **$6.40x10^{12}$** |
| **SD** | **0.7** | | | **$1.89x10^{12}$** |

**Table S6:** Hydrodynamic diameter and nanoparticle concentration for ferucarbotran. Values were corrected for dilution.

| Sample | Diameter (nm) | Attenuation Level (%) | Laser Power (%) | Particle Concentration (particles/mL) |
|---|---|---|---|---|
| Ferucarbotran | 67.7 | 75 | 80 | $2.39x10^{12}$ |
| Ferucarbotran | 65.9 | 75 | 100 | $2.64x10^{12}$ |
| Ferucarbotran | 68.8 | 75 | 100 | $2.03x10^{12}$ |
| **Average** | **67.5** | | | **$2.35x10^{12}$** |
| **SD** | **1.2** | | | **$2.53x10^{11}$** |

**Table S7:** Size and particle concentration for ABS as determined by NTA.

| Average Mean size (nm) | Average Mode size (nm) | Average Concentration, particles/mL |
|---|---|---|
| 99.1 | 76.6 | $2.47x10^{13}$ |

**Table S8:** Fe, Bi and S concentration in BSA, ABS and AB as determined by ICP-MS. Numbers represent mean values ± standard deviation.

| Sample | [Fe], mg/mL | [Bi], mg/mL | [S], mg/mL | # S/ BSA | [BSA], mg/mL | S:Bi ratio |
|---|---|---|---|---|---|---|
| BSA | N/A | N/A | 15.80 ± 1.52 (per g BSA) | 32 | N/A | N/A |
| ABS | 1.23 ± 0.05 | 5.09 ± 0.52 | 1.35 ± 0.08 | N/A | 84.6 ± 5 | 1.14 ± 0.06 |
| AB | -0.01 ± 0.72 | 4.85 ± 0.53 | 1.73 ± 0.08 | N/A | 108.5 ± 5 | 1.33 ± 0.05 |

**<u>Table S9:</u>** Particle concentration in ABS and AB as determined by single particle ICP-MS. Data represent three independent runs. Samples were diluted 500,000,000 (500M), 100,000,000 (100M) or 200,000,000 (200M) fold in water before analysis.

| | Sample | Counts | Mass/ part. Bi, (g) | Mean Dissolved Size (nm) | Mode Dissolved Size (nm) | SD Dissolved Size (nm) | Mass/ part. Bi2S3, (g) | Mean Dissolved Size (nm) ($Bi_2S_3$) | Mode Dissolved Size (nm) ($Bi_2S_3$) | SD Dissolved Size (nm) ($Bi_2S_3$) | Conc., part/mL | # Parts. | [Bi], mg/mL |
|---|---|---|---|---|---|---|---|---|---|---|---|---|---|
| Run #1 | ABS | 48.79 | $3.05x10^{-16}$ | 33.68 | 26.43 | 13.10 | $2.71x10^{-16}$ | 40.78 | 32.00 | 15.86 | $1.48x10^{13}$ | 793 | 4.53 |
| | ABS 500M | 49.08 | $3.07x10^{-16}$ | 34.04 | 27.79 | 13.02 | $2.72x10^{-16}$ | 41.21 | 33.64 | 15.76 | $1.48x10^{13}$ | 1621 | 4.56 |
| | AB | 28.99 | $1.81x10^{-16}$ | 29.01 | 24.32 | 9.79 | $1.60x10^{-16}$ | 35.12 | 29.45 | 11.85 | $2.48x10^{13}$ | 1396 | 4.48 |
| | AB 500M | 28.25 | $1.76x10^{-16}$ | 29.23 | 24.23 | 9.60 | $1.56x10^{-16}$ | 35.39 | 29.34 | 11.63 | $2.33x10^{13}$ | 2664 | 4.10 |
| Run #2 | ABS 100M | 31.84 | $2.57x10^{-16}$ | 33.08 | 28.22 | 11.14 | $2.28x10^{-16}$ | 40.05 | 34.16 | 13.49 | $2.04x10^{13}$ | 5535 | 5.24 |
| | ABS | 35.45 | $2.87x10^{-16}$ | 33.52 | 28.10 | 11.88 | $2.55x10^{-16}$ | 40.58 | 34.02 | 14.38 | $1.69x10^{13}$ | 931 | 4.85 |
| | ABS 500M | 34.67 | $2.81x10^{-16}$ | 33.38 | 27.46 | 11.79 | $2.49x10^{-16}$ | 40.41 | 33.24 | 14.27 | $1.94x10^{13}$ | 2094 | 5.45 |
| Run #3 | ABS 100M | 54.93 | $2.96x10^{-16}$ | 33.45 | 25.65 | 13.38 | $2.62x10^{-16}$ | 40.50 | 31.05 | 16.20 | $1.67x10^{13}$ | 7208 | 4.95 |
| | ABS | 56.18 | $3.03x10^{-16}$ | 33.40 | 26.50 | 13.54 | $2.68x10^{-16}$ | 40.43 | 32.08 | 16.39 | $1.64x10^{13}$ | 949 | 4.95 |
| | ABS 200M | 55.72 | $3.00x10^{-16}$ | 33.61 | 25.60 | 13.45 | $2.66x10^{-16}$ | 40.68 | 31.00 | 16.29 | $1.65x10^{13}$ | 3545 | 4.96 |
| | ABS 500M | 52.69 | $2.84x10^{-16}$ | 33.16 | 24.25 | 13.02 | $2.51x10^{-16}$ | 40.15 | 29.36 | 15.76 | $1.77x10^{13}$ | 1481 | 5.03 |
| | AB 100M | 54.38 | $2.93x10^{-16}$ | 33.05 | 24.25 | 13.50 | $2.60x10^{-16}$ | 40.01 | 29.36 | 16.34 | $1.78x10^{13}$ | 7431 | 5.21 |
| | AB | 49.68 | $2.67x10^{-16}$ | 32.11 | 25.53 | 13.01 | $2.37x10^{-16}$ | 38.88 | 30.91 | 15.75 | $1.80x10^{13}$ | 707 | 4.82 |
| | AB 200M | 51.96 | $2.80x10^{-16}$ | 32.71 | 24.96 | 13.18 | $2.48x10^{-16}$ | 39.60 | 30.23 | 15.95 | $1.73x10^{13}$ | 3589 | 4.84 |
| | AB 500M | 61.34 | $3.31x10^{-16}$ | 34.17 | 25.92 | 13.40 | $2.94x10^{-16}$ | 41.37 | 31.38 | 16.23 | $1.70x10^{13}$ | 1407 | 5.62 |
| **Average** | | | | | | | | | | | | | |
| | ABS | 48.28 | $2.96x10^{-16}$ | 33.50 | 26.61 | 12.92 | $2.63x10^{-16}$ | 40.56 | 32.22 | 15.64 | $1.72x10^{13}$ | 24313 | 5.09 |
| | AB | 45.77 | $2.55x10^{-16}$ | 31.71 | 24.87 | 12.08 | $2.26x10^{-16}$ | 38.40 | 30.11 | 14.63 | $1.97x10^{13}$ | 17194 | 4.84 |

**Table S10:** Particle concentration in ABS and AB as determined by DLS, NTA, and single particle ICP-MS.

| | Light Scattering | NTA | spICP-MS |
|---|---|---|---|
| ABS | $2.90 \times 10^{12}$ | $2.47 \times 10^{13}$ | $1.72 \times 10^{13}$ |
| AB | $6.40 \times 10^{12}$ | NT | $1.97 \times 10^{13}$ |

NT=not tested

**Table S11:** Zeta potential for ABS, AB, and ferucarbotran.

| Sample | pH | Zeta Potential mV |
|---|---|---|
| ABS | 7.29 (native) | -23.1 ± 1.5 |
| AB | 9.25 (native) | -24.3 ± 1.0 |
| AB | 7.59 (adjusted) | -20.2 ± 1.8 |
| Ferucarbotran | 7.42 (native) | -3.3 ± 0.7 |
